# Xrp1 and Irbp18 trigger a feed-forward loop of proteotoxic stress to induce the loser status

**DOI:** 10.1101/2021.07.02.450835

**Authors:** Paul F. Langton, Michael E. Baumgartner, Remi Logeay, Eugenia Piddini

## Abstract

Cell competition induces the elimination of less-fit “loser” cells by fitter “winner” cells. In *Drosophila*, cells heterozygous mutant in ribosome genes, *Rp/+*, known as Minutes, are eliminated via cell competition by wild-type cells. *Rp/+* cells display proteotoxic stress and the oxidative stress response, which drive the loser status. Minute cell competition also relies on the activities of the transcription factors Irbp18 and Xrp1, however how these contribute to the loser status is partially understood. Here, we show that Irbp18 and Xrp1 induce the loser status by promoting proteotoxic stress. We find that Xrp1 is necessary for *Rp/+* - induced proteotoxic stress and is sufficient to induce proteotoxic stress in otherwise wild-type cells. Xrp1 is also induced downstream of proteotoxic stress and required for the competitive elimination of cells suffering from proteotoxic stress. Our data suggests that a feed-forward loop between Xrp1, proteotoxic stress, and Nrf2 drives Minute cells to become losers.

## Introduction

Cells within a tissue may become damaged due to spontaneous or environmentally induced mutations, and it is beneficial to organismal health if these cells are removed and replaced by healthy cells. During cell competition, fitter cells, termed winners, recognise and eliminate less-fit cells, termed losers, resulting in restoration of tissue homoeostasis (Amoyel & Bach, 2014; Baker, 2011; Maruyama & Fujita, 2017). Cell competition therefore promotes tissue health and is thought to provide a level of protection against developmental aberrations (Baillon & Basler, 2014; Baker, 2017; Vincent, Kolahgar, Gagliardi, & Piddini, 2011) and against cancer by removing cells carrying oncoplastic mutations (Maruyama & Fujita, 2017; Vishwakarma & Piddini, 2020). However, an increasing body of evidence indicates that cell competition can also promote growth of established tumours, enabling them to expand at the expense of surrounding healthy cells (Vishwakarma & Piddini, 2020).

Minute cell competition was discovered through the study of a class of *Drosophila* ribosomal mutations called *Minutes* (Morata & Ripoll, 1975) and initial work suggests that it is conserved in mammals (Oliver, Saunders, Tarle, & Glaser, 2004). While homozygous *Rp* mutations are mostly cell lethal, heterozygosity for most *Rp* mutations gives rise to viable adult flies that exhibit a range of phenotypes including developmental delay and shortened macrochaete bristles (Marygold et al., 2007; Morata & Ripoll, 1975). *Rp/+* tissues display a higher cell-autonomous death frequency than wild-type tissues (Akai, Ohsawa, Sando, & Igaki, 2021; Baumgartner, Dinan, Langton, Kucinski, & Piddini, 2021; Coelho et al., 2005; Recasens-Alvarez et al., 2021). Competitive interactions further elevate cell death in *Rp/+* cells bordering wild-type cells, contributing to progressive loss of *Rp/+* clones over time (Baker, 2020; Baumgartner et al., 2021).

It was suggested that *Rp/+* cells are eliminated by cell competition due to their reduced translation rate (Amoyel & Bach, 2014; Kale et al., 2018; Lee et al., 2018; Milan, 2002; Moreno & Basler, 2004; Nagata, Nakamura, Sanaki, & Igaki, 2019). However, we and others have recently shown that *Rp/+* cells experience significant proteotoxic stress and this is the main driver of their loser status (Baumgartner et al., 2021; Recasens-Alvarez et al., 2021). *Rp/+* cells have a stochiometric imbalance of ribosome subunits, which may provide the source of proteotoxic stress. The autophagy and proteasomal machineries become overloaded and protein aggregates build up in *Rp/+* cells, leading to activation of stress pathways. This includes activation of Nuclear factor erythroid 2-related factor 2 (Nrf2) and of the oxidative stress response (Ma, 2013), which we have shown to be sufficient to cause the loser status (Kucinski, Dinan, Kolahgar, & Piddini, 2017). Restoring proteostasis in *Rp/+* cells suppresses the activation of the oxidative stress response and inhibits both autonomous and competitive cell death (Baumgartner et al., 2021; Recasens-Alvarez et al., 2021).

Genetic screening for suppressors of cell competition led to the identification of Xrp1 (Baillon, Germani, Rockel, Hilchenbach, & Basler, 2018; Lee et al., 2018), a basic leucine Zipper (bZip) transcription factor. Loss of Xrp1 rescues both the reduced growth and competitive cell death of *Rp/+* clones in mosaic tissues. Consistently, loss of Xrp1 restores translation rates and abolishes the increased JNK pathway activity characteristic of *Rp/+* cells (Lee et al., 2018). Xrp1 forms heterodimers with another bZip transcription factor called Inverted repeat binding protein 18kDa (Irbp18) (Francis et al., 2016; Reinke, Baek, Ashenberg, & Keating, 2013), and removal of Irbp18 also strongly suppresses the competitive elimination of *Rp/+* clones in mosaic tissues (Blanco, Cooper, & Baker, 2020). Irbp18 and Xrp1 are transcriptionally upregulated and mutually required for each other’s expression in *Rp/+* cells, suggesting they function together in Minute cell competition (Blanco et al., 2020). Irbp18 forms heterodimers with another bZip transcription factor, ATF4 (Reinke et al., 2013). Knockdown of ATF4 in *Rp/+* cells reduces their survival in mosaic tissues, which is the opposite effect to knockdown of Xrp1 or Irbp18. This has been interpreted to suggest that the ATF4-Irbp18 heterodimer acts independently to the Xrp1-Irbp18 heterodimer (Blanco et al., 2020).

How Xrp1/Irbp18 contribute to the loser status is not clear. Given the recently identified role of proteotoxic stress in cell competition we sought to establish whether Xrp1/Irbp18 and proteotoxic stress act in a linear pathway or independently contribute to cell competition in *Rp/+* cells. We identify a feed-forward loop between Xrp1/Irbp18 and proteotoxic stress, which is required for downstream activation of the oxidative stress response and the loser status. We find that the initial insult in *Rp/+* cells is ribosomal imbalance-induced proteotoxic stress. Xrp1 is transcriptionally activated downstream of proteotoxic stress both by the Unfolded Protein Response (UPR) and by Nrf2. The Xrp1-Irbp18 complex then induces further proteotoxic stress, completing the feed-forward loop. This work provides new insight into the interactions between the stress signalling pathways active in *Rp/+* cells and provides a mechanism for how the Xrp1-Irbp18 heterodimer mediates the competitive elimination of *Rp/+* cells by wild-type cells.

## Results

To probe the role of the Xrp1-Irbp18 complex in *Rp/+* cells, we first established whether RNAi lines against each functionally knock-down these genes. *xrp1* expression depends on its own activity and on the activity of Irbp18 (Blanco et al., 2020). As expected, knockdown of Xrp1 (*xrp1^KK104477^* RNAi line, hereafter referred to as *xrp1-RNAi*) in the posterior compartment of wild type wing discs reduced expression of an *xrp1* transcriptional reporter, *xrp1-lacZ* (Figure S1a-b). Similarly, knockdown of Irbp18 (*irbp18^KK110056^* RNAi line, hereafter referred to as *irbp18-RNAi*) reduced levels of *xrp1-lacZ* (Figure S1c-d). Mutations in *xrp1* and *irbp18* prevent *Rp/+* cells from being out-competed by wild-type cells in mosaic tissues (Baillon et al., 2018; Blanco et al., 2020; Lee et al., 2018). Accordingly, knockdown of Xrp1 or Irbp18 rescued the competitive elimination of *Rp/+* cells in wing discs. Compared to *Rp/+* clones, *Rp/+* clones expressing *xrp1-RNAi* (Figure S1e-g), or *irbp18-RNAi* (Figure S1h-j) grew substantially larger. These data indicate that those RNAi lines effectively knockdown Xrp1 and Irbp18.

To investigate the role of Xrp1 and Irbp18 in proteotoxic stress and the oxidative stress response, which are primary drivers of the loser status in *Rp/+* cells (Baumgartner et al., 2021; Kucinski et al., 2017; Recasens-Alvarez et al., 2021), we expressed *xrp1-RNAi* specifically in the posterior compartment of *Rp/+* wing discs with the *hedgehog (hh)-gal4* driver. Xrp1 knockdown significantly rescued the accumulation of phosphorylated-eukaryotic Initiation Factor 2α (p-eIF2α), a marker of the integrated stress response, which is induced in response to proteotoxic stress (Cnop, Toivonen, Igoillo-Esteve, & Salpea, 2017; Hetz, 2012), and is upregulated in *Rp/+* cells (Figure 1a-b). Xrp1 knockdown also strongly inhibited the oxidative stress response in *Rp/+* cells, as it reduced the expression of Glutathione S transferase D1-GFP (GstD1-GFP), a reporter of Nrf2 (Sykiotis & Bohmann, 2008) (Figure 1a and c). Irbp18 knockdown also rescued both p-eIF2α upregulation and GstD1-GFP upregulation in *Rp/+* discs (Figure 1d-f). Refractory to sigma P (Ref(2)p), also known as p62, is an autophagy adaptor and cargo (Mauvezin, Ayala, Braden, Kim, & Neufeld, 2014) and a marker of cytosolic protein aggregates (Nezis et al., 2008), which accumulates in *Rp/+* cells due to proteotoxic stress overload (Baumgartner et al., 2021). The accumulation of p62-labelled aggregates in *Rp/+* cells was rescued both by *xrp-RNAi* (Figure 1g-h) and *irbp18-RNAi* (Figure 1i-j), further indicating that proteotoxic stress in *Rp/+* cells is mediated by the Xrp1/Irbp18 complex. Together, these data show that Xrp1 and Irbp18 are required for, and act upstream of, proteotoxic stress and the oxidative stress response in *Rp/+* cells.

**Figure 1.**
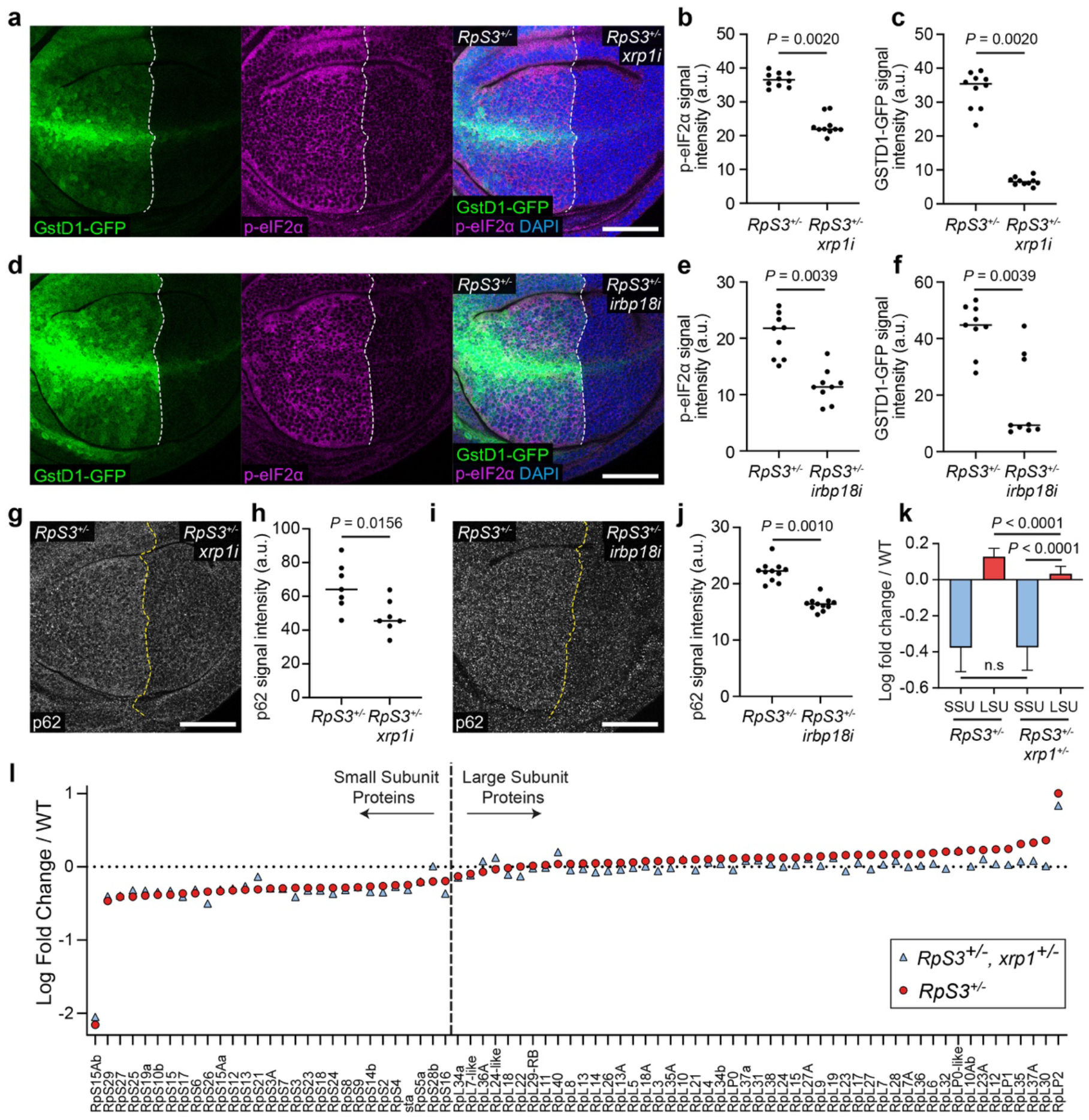
Xrp1 and Irbp18 are required for proteotoxic stress and the oxidative stress response induced by Rp loss. (**a-c**) An *RpS3^+/−^* wing disc harboring the GSTD1-GFP reporter (green) and expressing *xrp1-RNAi* (*xrp1i*) in the posterior compartment, immuno-stained for p-eIF2α (magenta) with nuclei labelled in blue (**a**). Quantifications of p-eIF2α signal intensity (n = 10; two-sided Wilcoxon signed-rank test) and GSTD1-GFP signal intensity (n = 10; two-sided Wilcoxon signed-rank test) are shown in (**b**) and (**c**) respectively. (**d-f**) An *RpS3^+/−^* wing disc harboring the GSTD1-GFP reporter (green) and expressing *irbp18-RNAi* (*irbp18i*) in the posterior compartment, immuno-stained for p-eIF2α (magenta) with nuclei labelled in blue (**d**). Quantifications of p-eIF2α signal intensity (n = 9; two-sided Wilcoxon signed-rank test) and GSTD1-GFP signal intensity (n = 9; two-sided Wilcoxon signed-rank test) are shown in (**e**) and (**f**) respectively. (**g-h**) A wing disc of the same genotype as shown in (**a**), immuno-stained for p62 (grey) (**g**), with quantification of p62 signal intensity (**h**) (n = 7; two-sided Wilcoxon signed-rank test). (**i-j**) A wing disc of the same genotype as shown in (**b**), immuno-stained for p62 (grey) (**i**), with quantification of p62 signal intensity (**j**) (n = 11; two-sided Wilcoxon signed-rank test). (**k**) A bar graph showing the mean log fold change in all Small-subunit (SSU) and Large-subunit (LSU) ribosomal proteins detected by mass spectrometry in *RpS3^+/−^* and *RpS3^+/−^, Xrp1^+/−^* wing discs relative to wild-type discs, as indicated (n = 29; two-sided Wilcoxon signed-rank test for comparison of SSU, n = 49; two-sided Wilcoxon signed-rank test for comparison of LSU, n = 29 and 49, respectively; two-sided Mann–Whitney U-test for comparison of SSU and LSU in *RpS3^+/−^, Xrp1^+/−^* wing discs), error bars represent 95% confidence interval. (**l**) Mean log fold change in SSU and LSU ribosomal proteins detected by mass spectrometry (n = 2) in *RpS3^+/−^* and *RpS3^+/−^, Xrp1^+/−^* wing discs relative to wild-type discs, as indicated. In this figure and throughout: scale bars are 50μm; dashed white or yellow lines mark compartment boundaries; each data point on the scatter plots represents one wing disc or one wing disc compartment and the horizontal line represents the median; all n values refer to the number of individual wing discs except for Figure 1k-l; posterior is right and dorsal is up.

Mutations in the E3 ubiquitin ligase encoding gene *mahjong* (*mahj)* lead to the loser status, and *mahj^−/−^* cells are out-competed by wild-type cells in mosaic tissues (Tamori et al., 2010). Although Mahj is functionally distinct to ribosomal proteins, the gene expression signatures of *mahj* and *RpS3* mutants significantly overlap (Kucinski et al., 2017), indicating a common mechanism leading to the loser status. Indeed, *mahj* cells also show upregulation of markers of proteotoxic stress (Baumgartner et al., 2021). Interestingly, Xrp1 knockdown rescued *mahj*-RNAi expressing clones from elimination in mosaic wing discs (Figure S1k-m). Thus, Xrp1 contributes to the competitive elimination of cells deficient in two distinct loser mutations, *Rp/+* and *mahj*, which are both linked to proteotoxic stress.

*Rp/+* cells have recently been shown to have a stochiometric imbalance in their ribosome subunits, suggesting that this is the initial proteostatic perturbation leading to proteotoxic stress. Specifically, *Rp/+* cells have an excess of large-subunit (LSU) proteins and a reduced complement of small-subunit (SSU) proteins, relative to wild-type cells (Baumgartner et al., 2021; Recasens-Alvarez et al., 2021). The data in Figure 1a-j indicate that proteotoxic stress is induced by Xrp1 and Irbp18 in *Rp/+* cells, therefore we asked whether the ribosomal imbalance in *Rp/+* cells is also downstream of Xrp1. Interestingly, proteomic analysis revealed that removal of one copy of *xrp1*, which is sufficient to rescue *Rp/+* cells from competition (Lee et al., 2018), rescues the excess of LSU proteins but does not affect the reduction in SSU proteins (Figure 1k-l). Thus, SSU protein imbalance is independent of Xrp1. This suggests that the initial proteotoxic stress experienced by *Rp/+* cells is an SSU/LSU stoichiometric imbalance. This may provide the signal for Xrp1 induction, which in turn exacerbates proteotoxic stress, resulting in accumulation of LSU proteins.

The results described above suggest that Xrp1 functions upstream of proteotoxic stress in *Rp/+* cells, so we asked whether Xrp1 is sufficient to induce proteotoxic stress. We over-expressed the *xrp1^long^* isoform (Tsurui-Nishimura et al., 2013) in the posterior compartment of wing discs with the *engrailed (en)-gal4* driver and found this condition to be larval lethal before the 3^rd^ instar, which is consistent with previous reports that *xrp1* over-expression induces high levels of cell death (Blanco et al., 2020; Boulan, Andersen, Colombani, Boone, & Leopold, 2019; Tsurui-Nishimura et al., 2013). To circumvent this lethality, we used a temperature sensitive Gal4 inhibitor, Gal80^ts^, to prevent *xrp1* expression throughout most of larval development. Shifting the larvae to the Gal80^ts^ restrictive temperature 24 hours before dissection allowed for a relatively short burst of *xrp1* expression. Under these conditions, *xrp1* over-expressing compartments accumulated p62 (Figure 2a-b) and had higher levels of p-eIF2α (Figure 2c-d) than the wild-type, control compartments. Therefore, Xrp1 is sufficient to induce proteotoxic stress. Furthermore, *xrp1* expression led to a strong increase in GstD1-GFP levels (Figure 2e-f), indicating that the oxidative stress response is also activated downstream of Xrp1. Overall, this data suggests that Xrp1 and Irbp18 are responsible for inducing proteotoxic stress and the oxidative stress response in *Rp/+* cells, which explains why their removal so effectively rescues Minute competition.

**Figure 2.**
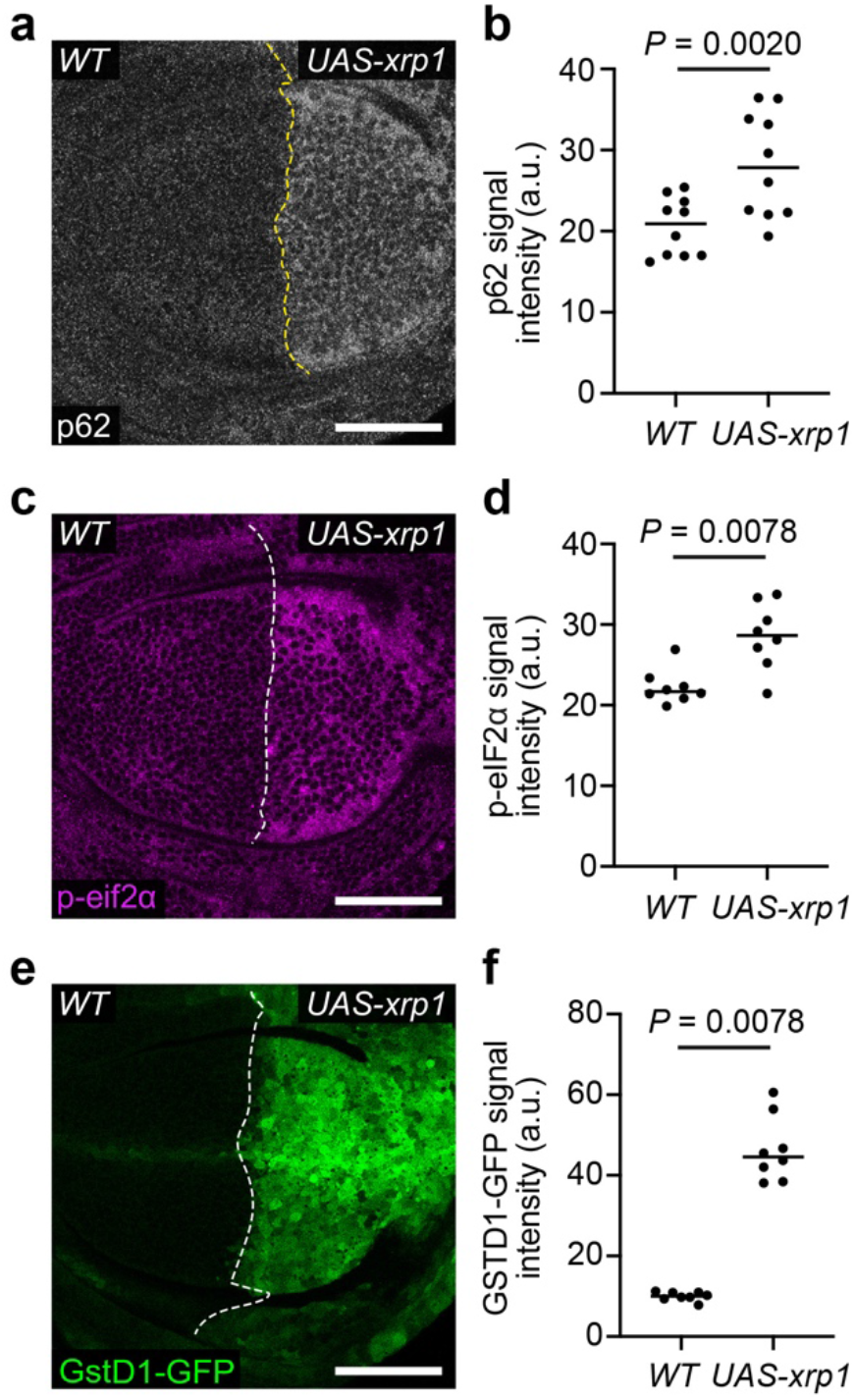
Xrp1 is sufficient for proteotoxic stress and the oxidative stress response. (**a-b**) A wild-type (WT) wing disc harboring GSTD1-GFP and over-expressing *xrp1* (*UAS-xrp1*) in the posterior compartment and immuno-stained for p62 (grey) (**a**) with quantification of p62 signal intensity (**b**) (n = 10; two-sided Wilcoxon signed-rank test). (**c-d**) A wing disc of the same genotype as in (**a**) immuno-stained for p-eIF2α (magenta) (**c**) with quantification of p-eIF2α signal intensity (**d**) (n = 8; two-sided Wilcoxon signed-rank test). (**e-f**) GSTD1-GFP reporter signal (green) (**e**) in a wing disc of the same genotype as in (**a**) with quantification of GSTD1-GFP signal intensity shown in (**f**) (n = 8; two-sided Wilcoxon signed-rank test).

If Xrp1 and Irbp18 are required in competition because they induce proteotoxic stress, then inducing proteotoxic stress by other means should lead to the loser status in an Xrp1- and Irbp18-independent manner. To test this hypothesis, we induced proteotoxic stress by well-established means. eIF2α is phosphorylated in response to proteotoxic stress, leading to global attenuation of translation (Cnop et al., 2017; Hetz, 2012). However, sustained increase in p-eIF2α has also been shown to induce proteotoxic stress, by causing accumulation of aggregogenic stress granules (Baradaran-Heravi, Van Broeckhoven, & van der Zee, 2020; Ohno, 2014). Therefore, we sought to induce high levels of p-eIF2α. Growth arrest and DNA-damage-inducible 34 (GADD34) is a Protein Phosphatase 1 (PP1) regulatory subunit, which causes p-eIF2α dephosphorylation by providing PP1 with target specificity for p-eIF2α (Novoa, Zeng, Harding, & Ron, 2001). As expected, *GADD34-RNAi* increased the levels of p-eIF2α (Figure S2a-b). *GADD34-RNAi* in the posterior compartment of wing discs also led to higher levels of p62 (Figure 3a-b) and of mono- and poly-ubiquitinated proteins (detected by the FK2 antibody) than in the control anterior compartment (Figure 3c-d). As these are both markers of protein aggregates (Nezis et al., 2008; Rubinsztein, 2006), these data indicate that sustained eIF2α phosphorylation induces proteotoxic stress and protein aggregation. GADD34 knockdown also upregulated GstD1-GFP (Figure 3e-f) and p-JNK (Figure 3g and Figure S2c). Thus, increased levels of p-eIF2α are sufficient to induce proteotoxic stress, the oxidative stress response, and JNK pathway activity, all of which are observed in *Rp/+* prospective loser cells.

**Figure 3.**
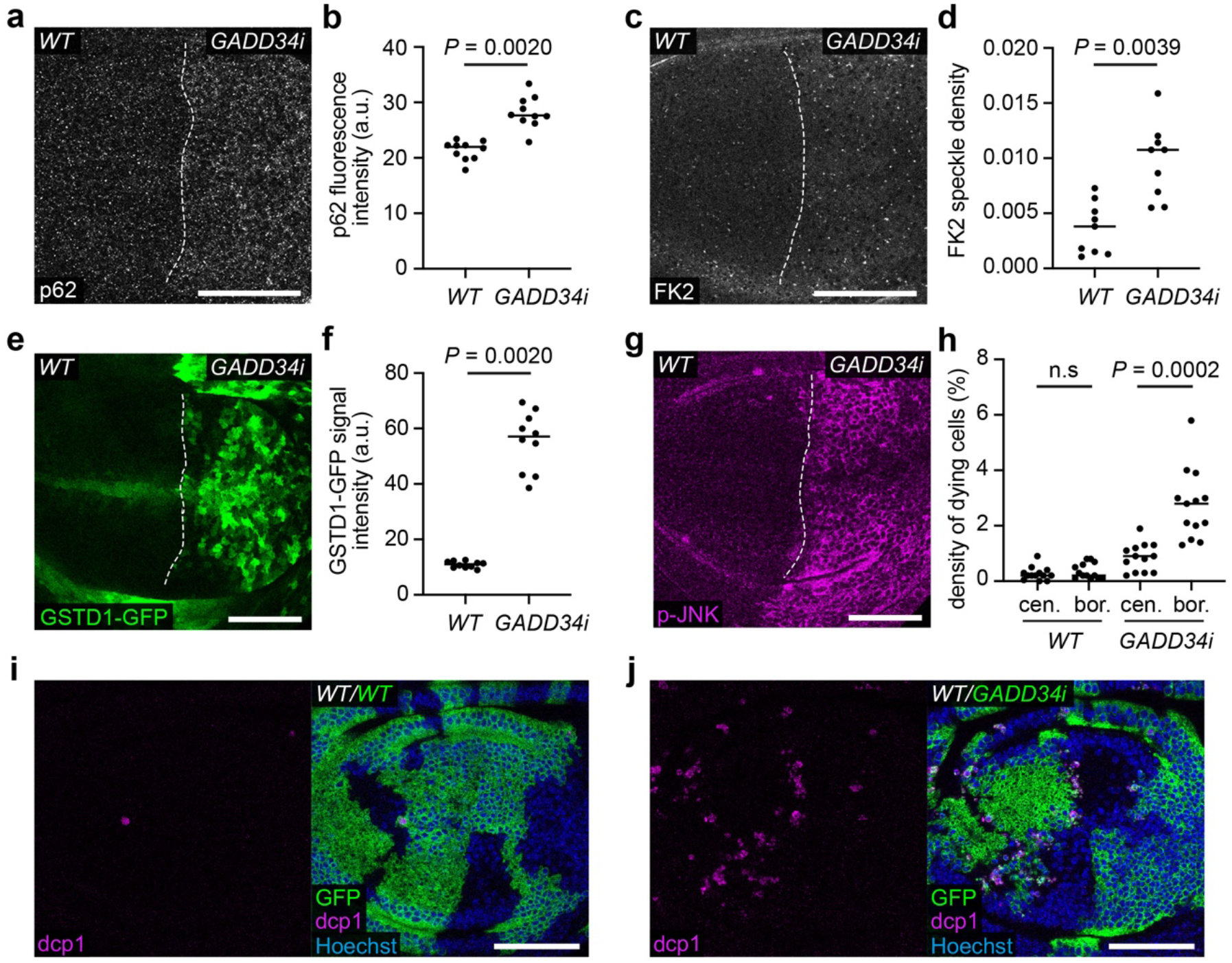
GADD34 knockdown induces proteotoxic stress and the loser status. (**a-b**) A wild-type (WT) wing disc carrying the GSTD1-GFP reporter and expressing *GADD34-RNAi* (*GADD34i*) in the posterior compartment, immuno-stained for p62 (grey) (**a**) with quantification of p62 fluorescence intensity (**b**) (n = 10; two-sided Wilcoxon signed-rank test). (**c-d**) A wing disc of the same genotype as in (**a**), immuno-stained for FK2 (grey) to label mono- and poly-ubiquitinated proteins (**c**) with quantification of FK2 speckle density (**d**) (n = 9; two-sided Wilcoxon signed-rank test). (**e-f**) GSTD1-GFP (green) in a wing disc of the same genotype as in (**a**), with quantification of GSTD1-GFP signal intensity (**f**) (n = 10; two-sided Wilcoxon signed-rank test). (**g-h**) A wing disc of the same genotype as in (**a**), immuno-stained for p-JNK (magenta) (**g**). (**h-j**) Wing discs harboring either WT clones (GFP positive) (**i**) or *GADD34-RNAi* expressing clones (GFP positive) (**j**) immuno-stained for dcp1 (magenta), with quantification of density of dying cells at the center (cen.) and border (bor.) of clones as indicated (**h**) (n = 13 and 13, respectively; two-sided Wilcoxon signed-rank test). The clone border defines cells within two cell diameters of the clone perimeter.

We then induced *GADD34-RNAi* in a mosaic fashion to test whether it induces the loser status. *GADD34-RNAi* expressing clones were efficiently removed from wing discs in mosaic experiments (Figure S2d). Only a few fragments of clones remained, and these had been basally extruded from the epithelium (Figure S2e), consistent with competitive elimination. However, it was also possible that this was due to cell-autonomous activation of apoptosis. Thus, we designed an experimental strategy to obtain large clones (Figure S2f) and directly compare the rate of apoptosis at clone borders and centers, as increased border death is a hallmark of cell competition (Baker, 2020; Li & Baker, 2007). We made use of Gal80^ts^ for conditional expression and placed larvae at the Gal80^ts^ permissive temperature after clone induction, to allow clones to expand without induction of transgene expression. We then induced *GADD34-RNAi* (and *GFP*) expression by moving larvae to the Gal80^ts^ restrictive temperature 24 hours before dissection (Figure S2f). This short period of *GADD34-RNAi* expression was sufficient to increase p-eIF2α (Figure S2g). Unlike control wild-type clones, *GADD34-RNAi* expressing clones had significantly higher levels of cell death at clone borders than in the center of clones, showing that they are subject to competitive elimination by wild-type cells (Figure 3h-j).

We next asked whether *GADD34-RNAi* induced cell elimination depends on Xrp1. As Xrp1 and Irbp18 function upstream of proteotoxic stress in *Minutes* (Figure 1), we were surprised to find that co-expression of *xrp1-RNAi* with *GADD34-RNAi* resulted in a strong rescue of clone elimination (Figure 4a-c). Clones were readily recovered in every disc, suggesting that Xrp1 also functions downstream of proteotoxic stress. Formally, one possible explanation for this rescue could be the presence of a second UAS construct (*UAS-xrp1-RNAi)*, which, by titrating Gal4, could weaken the expression of *UAS-GADD34-RNAi*. To rule out this possibility, the clones expressing *GADD34-RNAi* alone in this experiment also carried a second UAS construct: an inert UAS insertion on the second chromosome that doesn’t drive expression of a transgene. All further experiments in this study that compare the phenotype of expression of a single UAS construct to that of two UAS constructs use this strategy. Thus, *GADD34-RNAi* clone elimination is mediated by Xrp1. Altogether, these data show that Xrp1 functions both upstream and downstream of proteotoxic stress and suggest that a feed-forward loop between proteotoxic stress and Xrp1 drives *Rp/+* cells to become losers.

**Figure 4.**
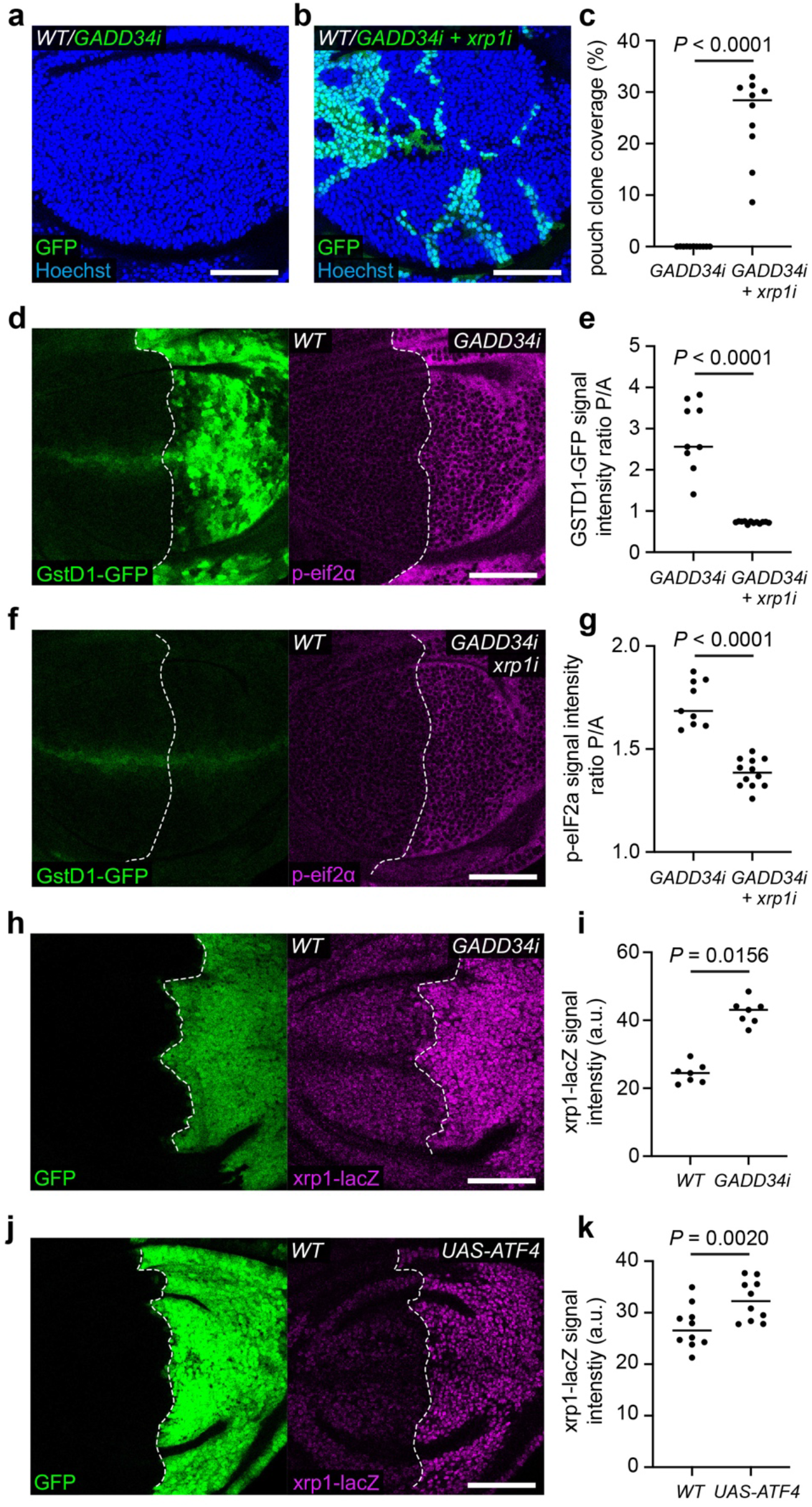
A feed-forward loop between Xrp1 and proteotoxic stress. (**a-b**) Wild-type wing discs harboring *GADD34-RNAi* (*GADD34i*) expressing clones (GFP positive) (**a**) or *GADD34-RNAi* and *xrp1-RNAi* (*xrp1i*) expressing clones (GFP positive) (**b**) with nuclei labelled in blue, and quantification of percentage clone coverage of the pouch (**c**) (n = 11 and 10, respectively; two-sided Mann–Whitney U-test). (**d-g**) Wing discs harboring GSTD1-GFP (green) and expressing either *GADD34-RNAi* (**d**) or *GADD34-RNAi* and *xrp1-RNAi* (**f**) in the posterior compartment, immuno-stained for p-eIF2α (magenta), with quantification of the Posterior / Anterior (P/A) ratio of GSTD1-GFP signal intensity (**e**) (n = 9 and 12, respectively; two-sided Student’s t-test) and the Posterior / Anterior (P/A) ratio of p-eIF2α signal intensity (**g**) (n = 9 and 12, respectively; two-sided Student’s t-test). (**h-i**) A wing disc carrying the *xrp1-lacZ* reporter and expressing *GADD34-RNAi* and *GFP* (green) in the posterior compartment, immuno-stained with anti-β-galactosidase (magenta) (**h**), with quantification of *xrp1-lacZ* signal intensity (**i**) (n = 7; two-sided Wilcoxon signed-rank test). (**j-k**) A wing disc carrying the *xrp1-lacZ* reporter and over-expressing *ATF4* (*UAS-ATF4*) and *GFP* (green) in the posterior compartment, immuno-stained with anti-β-galactosidase (magenta) (**j**), with quantification of *xrp1-lacZ* signal intensity (**k**) (n = 10; two-sided Wilcoxon signed-rank test).

Xrp1 knockdown completely rescued the increased GstD1-GFP observed in *GADD34-RNAi* expressing compartments, bringing levels down to, or even slightly lower than, wild-type levels (Figure 4d-f). Remarkably, Xrp1 knockdown was also able to partially rescue the increased p-eIF2α in *GADD34-RNAi* expressing compartments (Figure 4d and f-g), suggesting that removing Xrp1 breaks the feed-forward loop to proteotoxic stress, and therefore partially rescues the increased p-eIF2α in *GADD34-RNAi* expressing cells.

How might Xrp1 be acting downstream of proteotoxic stress? During ER stress, the UPR induces eIF2α phosphorylation, which mediates global translation repression and selective translation of a subset of transcripts, including that of ATF4, which, in mammals, mediates expression of chaperones and proapoptotic genes, including *CHOP* (Cnop et al., 2017; Hetz, 2012). Although no clear mammalian Xrp1 homologs exist, it has been suggested that Xrp1 is the functional homolog of CHOP (Blanco et al., 2020), which heterodimerizes with CEBPγ, the human homolog of Irbp18 (Deppmann, Alvania, & Taparowsky, 2006). Consistently, we found that *GADD34-RNAi* expressing compartments had significantly higher *xrp1-lacZ* signal than control compartments (Figure 4h-i). Furthermore, *ATF4* over-expressing compartments upregulated *xrp1-lacZ* (Figure 4j-k). These data therefore suggest that ATF4 mediates Xrp1 transcriptional activation in *GADD34-RNAi* and *Rp/+* cells, mirroring CHOP regulation by ATF4 during the UPR in mammals.

We have previously shown that proteotoxic stress induces expression of the Nrf2 reporter GstD1-GFP (Baumgartner et al., 2021) and that over-expression of *nrf2* is sufficient to turn otherwise wild-type cells into losers (Kucinski et al., 2017). Therefore, we investigated whether the contributions of Nrf2 and of Xrp1 to the loser status are functionally linked. In the absence of Gal80^ts^, *nrf2* expressing clones were readily eliminated from wing discs, with only a few tiny clones remaining at the time of dissection (Figure 5a and c). *xrp1-RNAi* significantly rescued the growth of *nrf2* expressing clones (Figure 5b-c), indicating that Xrp1 functions downstream of Nrf2. Irbp18 knockdown also rescued *nrf2* expressing clones from elimination (Figure 5d-f) confirming that Xrp1 functions along with Irbp18, downstream of Nrf2. This suggests that in *Rp/+* tissues, proteotoxic stress activates Xrp1 by two routes, one via the UPR and ATF4, and the other via Nrf2.

**Figure 5.**
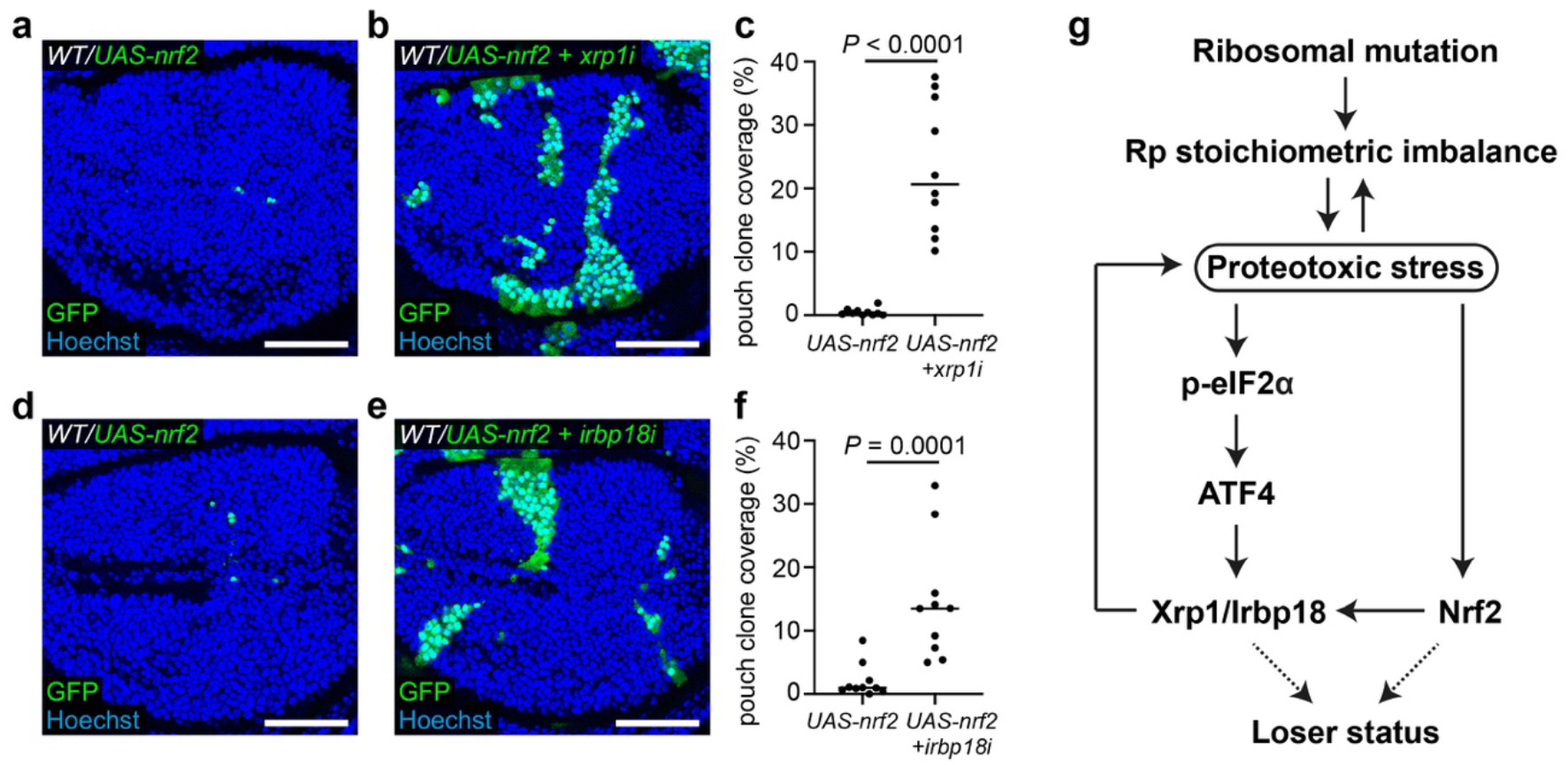
Xrp1 and Irbp18 function downstream of Nrf2. (**a-c**) Wild-type wing discs harboring *UAS-nrf2* expressing clones (GFP positive) (**a**) or *UAS-nrf2* and *xrp1-RNAi* (*xrp1i*) expressing clones (GFP positive) (**b**) with nuclei labelled in blue, and quantification of percentage clone coverage of the pouch (**c**) (n = 10 and 10, respectively; two-sided Mann– Whitney U-test). (**d-f**) Wild-type wing discs harboring *UAS-nrf2* expressing clones (GFP positive) (**d**) or *UAS-nrf2* and *irbp18-RNAi* (*irbp18i*) expressing clones (GFP positive) (**e**) with nuclei labelled in blue, and quantification of percentage clone coverage of the pouch (**f**) (n = 10 and 10, respectively; two-sided Mann–Whitney U-test). (**g**) Working model describing the role of the Xrp1-Irbp18 complex in *Rp/+* cells.

## Discussion

We have provided evidence for a feed-forward loop between proteotoxic stress and the Xrp1/Irbp18 complex, which is required for the elimination *Rp/+* cells in competing mosaic tissues (Figure 5g). Our data suggests that initial proteotoxic stress comes from an imbalance between SSU and LSU Ribosomal proteins. Proteotoxic stress in *Rp/+* cells induces Xrp1 expression both downstream of p-eIF2α, likely by the activity of ATF4, and possibly downstream of Nrf2. Xrp1 then acts, together with Irbp18, in a feed-forward loop, generating further proteotoxic stress. This causes LSU Rp’s to accumulate, exacerbating the stochiometric imbalance of Rp’s in *Rp/+* cells. Knockdown of Xrp1 and Irbp18 can rescue proteotoxic stress and the oxidative stress response in *Rp/+* cells, suggesting that this feed-forward loop is essential for build-up of proteotoxic stress and to reduce the competitiveness of *Rp/+* cells.

If ATF4 is partly responsible for Xrp1 upregulation in *Rp/+* cells, then removal of ATF4 would be expected to rescue Minute competition. However, the opposite effect has been observed, whereby *ATF4-RNAi* expressing clones generated in *Rp/+* discs are much smaller than control clones generated in *Rp/+* discs (Blanco et al., 2020), suggesting that ATF4 worsens, rather than rescues, the loser status. On the basis of our data, we favour a model whereby ATF4 plays a dual role: it promotes Xrp1 expression in *Rp/+* cells, possibly along with Irbp18; however, it is also required for expression of chaperones, which help cells cope with proteotoxic stress (Cnop et al., 2017; Hetz, 2012), explaining why removal of ATF4 worsens the loser status of *Rp/+* cells (Blanco et al., 2020). Of note, Nrf2 plays a similar dual role in *Minute* cell physiology; it contributes to the loser status, but is also cytoprotective in *Rp/+* cells (Kucinski et al., 2017), likely due to its ability to promote proteostasis (Baxter et al., 2021; Pajares, Cuadrado, & Rojo, 2017; Skibinski et al., 2017).

*CHOP* is one of several pro-apoptotic genes activated by the UPR to ensure that cells with extensive ER damage are eliminated (Cnop et al., 2017; Hetz, 2012; Hu, Tian, Ding, & Yu, 2018). *xrp1* expression is induced downstream of p-eIf2a and is required for the elimination of cells with high levels of p-eIF2a (Figure 4). Furthermore, Xrp1 overexpression induces high levels of cell death (Blanco et al., 2020; Tsurui-Nishimura et al., 2013), suggesting that, like CHOP, Xrp1 has a proapoptotic role in cells experiencing ER stress, which is consistent with the previously suggested notion that Xrp1 is functionally equivalent to CHOP (Blanco et al., 2020).

Ribosomopathies are a diverse class of human genetic disorders, resulting from either mutation of one copy of a Rp encoding gene or from defects in ribosome biogenesis (Mills & Green, 2017). A mutation equivalent to that of a ribosomopathy patient (*RpS23^R67K^*) was engineered in *Drosophila*, where it causes increased cell death and proteotoxic stress. Knockdown of Xrp1 rescues autonomous cell death in *RpS23^R67K^/+* cells (Recasens-Alvarez et al., 2021). Our work suggests that proteotoxic stress in *RpS23^R67K^/+* cells would also be rescued by removal of Xrp1 and, by extension, that CHOP, or regulators of CHOP, could be promising drug targets for therapeutics for ribosomopathies.

Nrf2 plays a pro-survival role in many contexts, by activating a battery of genes that enable the metabolic adaptation to oxidative stress (Ma, 2013). It is therefore counterintuitive that Nrf2 overexpression should induce the loser status and, at high expression levels, cell death (Kucinski et al., 2017). Our work suggests that the toxicity of Nrf2 is at least in part due to Xrp1 function as elimination of Nrf2 expressing cells is rescued by Xrp1 knockdown. Whether additional Nrf2 target genes contribute to the loser status remains to be established.

---

**Figure S1.**
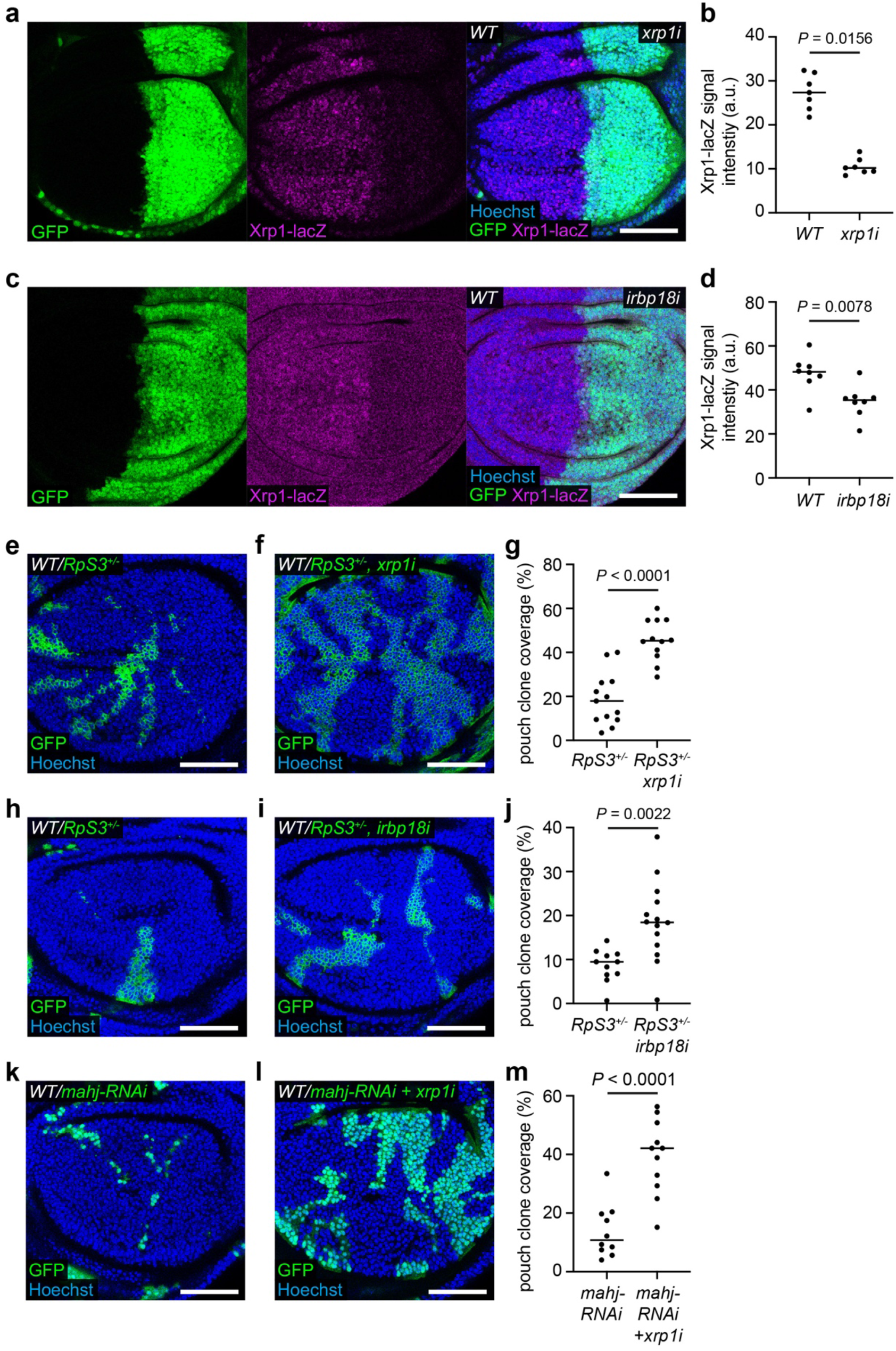
Xrp1 or Irbp18 knockdown reduces *xrp1* transcription and rescues elimination of *Rp/+* cells. (**a-b**) A wing disc carrying the *xrp1-lacZ* reporter and expressing *xrp1-RNAi* (*xrp1i*) and *GFP* (green) in the posterior compartment, immuno-stained with anti-β-galactosidase (magenta) and nuclei labelled with in blue (**a**), with quantification of *xrp1-lacZ* signal intensity (**b**) (n = 7; two-sided Wilcoxon signed-rank test). (**c-d**) A wing disc carrying the *xrp1-lacZ* reporter and expressing *irbp18-RNAi* (*irbp18i*) and *GFP* (green) in the posterior compartment, immuno-stained with anti-β-galactosidase (magenta) and nuclei labelled in blue (**c**), with quantification of *xrp1-lacZ* signal intensity (**d**) (n = 8; two-sided Wilcoxon signed-rank test). (**e-g**) Wild-type wing discs harboring *RpS3^+/−^* clones (GFP positive) (**e**) or *RpS3^+/−^* clones also expressing *xrp1-RNAi* (GFP positive) (**f**) with nuclei labelled in blue, and quantification of percentage clone coverage of the pouch (**g**) (n = 13 and 12, respectively; two-sided Student’s t-test). (**h-j**) Wild-type wing discs harboring *RpS3^+/−^* clones (GFP positive) (**h**) or *RpS3^+/−^* clones also expressing *irbp18-RNAi* (GFP positive) (**i**) with nuclei labelled in blue, and quantification of percentage clone coverage of the pouch (**j**) (n = 11 and 14, respectively; two-sided Student’s t-test). (**k-m**) Wild-type wing discs harboring *mahj-RNAi* clones (GFP positive) (**k**) or *mahj-RNAi* clones also expressing *xrp1-RNAi* (GFP positive) (**l**) with nuclei labelled in blue, and quantification of percentage clone coverage of the pouch (**m**) (n = 10 and 11, respectively; two-sided Student’s t-test).

**Figure S2.**
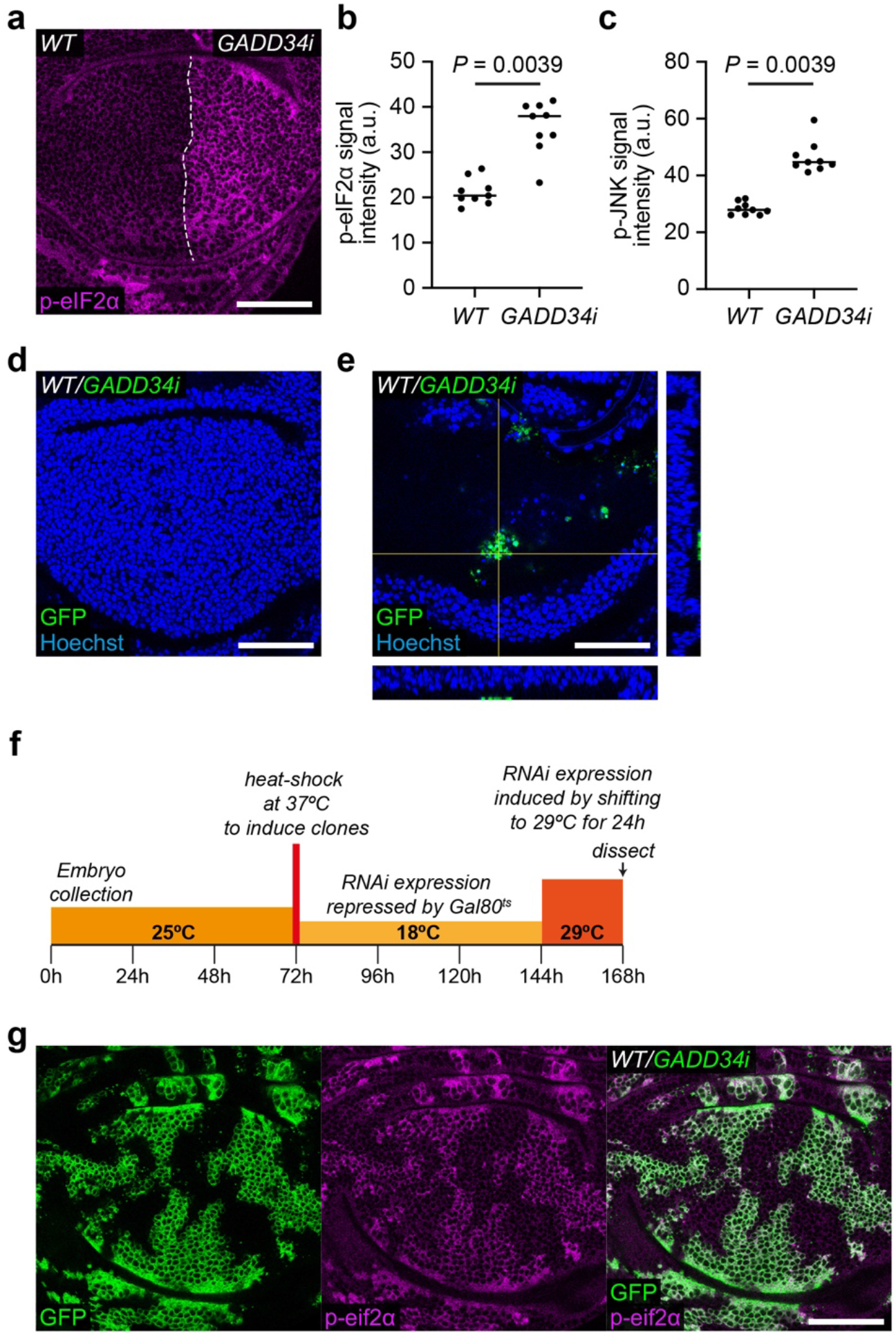
GADD34 knockdown induces the loser status. (**a-b**) A wing disc expressing *GADD34-RNAi* (*GADD34i*) in the posterior compartment and immuno-stained for p-eIF2α (magenta) (**a**) with quantification of p-eIF2α signal intensity (**b**) (n = 9; two-sided Wilcoxon signed-rank test). (**c**) Quantification of p-JNK signal intensity in wing discs expressing *GADD34-RNAi* in the posterior compartment (n = 9; two-sided Wilcoxon signed-rank test). (**d**) A wing disc harboring *GADD34-RNAi* expressing clones (GFP positive), generated in the absence of Gal80^ts^, with nuclei labelled in blue. (**e**) A basal section of a wing disc with *GADD34-RNAi* clones (GFP positive), generated in the absence of Gal80^ts^, with nuclei labelled in blue, to show that only small, basally extruded *GADD34-RNAi* expressing clones remain. Orthogonal views taken at the positions indicated by the yellow lines are shown to the right and bottom of the main image. (**f**) Schematic depicting experimental conditions for generating large *GADD34-RNAi* expressing clones. (**g**) A wing disc with *GADD34-RNAi* expressing clones (GFP positive), generated with the experimental conditions depicted in (**f**), immuno-stained for p-eIF2α (magenta).

## Acknowledgments

We thank members of the Piddini lab for helpful discussions on the project. We thank the Wolfson Bioimaging Facility for access to microscopes and the University of Bristol Proteomics Facility for performing the TMT proteomic experiments and for bioinformatics support. This work was supported a Cancer Research UK Programme Foundation Award to E.P. (Grant C38607/A26831) and a Wellcome Trust Senior Research Fellowship to E.P. (205010/Z/, 16/Z).

## Author contributions

E.P led the project. All authors conceived of the experiments. P.F.L performed and analysed the majority of the experiments with contributions from M.E.B and R.L. P.F.L and E.P wrote the manuscript with feedback from M.E.B and R.L.

## Financial and non-financial competing interests

The authors declare no competing interests.

## Materials & correspondence

The Lead Contacts, Professor Eugenia Piddini and Dr Paul F. Langton will fulfil requests for resources and reagents.

## Data availability

All source numerical data are provided in the Statistics Source Data table. All other data used in this paper are available upon reasonable request.

## Methods

### Fly husbandry

Fly food composition is: 7.5g/L agar powder, 50g/L baker’s yeast, 55g/L glucose, 35g/L wheat flour, 2.5 % nipagin, 0.4 % propionic acid and 1.0% penicillin/streptomycin. Eggs were collected for 24 hours in a 25°C incubator and experimental crosses were maintained in either an 18°C incubator, a 25°C incubator, or in a water bath set to a specific temperature as indicated in the genotypes table below. Wing discs were dissected from wandering third instar larvae. For all datasets, egg collections, heat shocks, temperature shifts, dissections, and imaging were done in parallel for control and experimental crosses. For mosaic competition experiments, all dissected larvae were of the same sex for both the control and experimental crosses. For half-half experiments, where the anterior compartment and posterior compartment were compared, sexes were not differentiated.

### Immunostaining

Wandering third instar larvae were dissected in phosphate buffered saline (PBS) and hemi-larvae were fixed in 4% paraformaldehyde for 20 minutes at room temperature. Tissues were permeabilized with three 10-minute washes in PBST (0.25% triton in PBS) and blocked for 20 minutes in blocking buffer (4% fetal calf serum in PBST). Samples were incubated with primary antibodies diluted in blocking buffer at the concentration indicated in the key resources table overnight at 4°C. Samples were washed three times in PBST for 10 minutes and incubated with secondary antibodies and Hoechst diluted in blocking buffer at the concentration indicated in the key resources table for 45-minutes at room temperature. After a further three 10-minute washes in PBST, wing discs were dissected from hemi-larvae and mounted in Vectashield (Vector laboratories) on borosilicate glass sides (no 1.5, VWR international).

### Clonal analysis and temperature shifts

Mosaic wing discs were generated with the hs-FLP transgenic line by heat shocking crosses three days after egg laying in a 37°C water bath for the time indicated in the genotypes table below. For experiments using temperature sensitive Gal80 (Gal80^ts^) to control the timing and level of transgene expression, conditions were optimized for each experiment and crosses were incubated in either an incubator or water bath set to the temperatures indicated in the genotypes table below.

Genotypes Table.

**Genotypes Table**
| Figure number/panel | Genotype | Experimental conditions |
| --- | --- | --- |
| Main figures |  |  |
| 1a | *GstD1-GFP/UAS-xrp1-RNAi; FRT82B, RpS3[Plac92], hh-Gal4/+* | 25°C |
| 1d | *GstD1-GFP/UAS-irbp18-RNAi; FRT82B, RpS3[Plac92], hh-Gal4/+* | 25°C |
| 1g | *GstD1-GFP/UAS-xrp1-RNAi; FRT82B, RpS3[Plac92], hh-Gal4/+* | 25°C |
| 1i | *GstD1-GFP/UAS-irbp18-RNAi; FRT82B, RpS3[Plac92], hh-Gal4/+* | 25°C |
| 1l (control) | *yw* | 25°C |
| 1l (*RpS3<sup>+/-</sup>*) | *FRT82B, RpS3[Plac92]/+* | 25°C |
| 1l (*RpS3<sup>+/-</sup>, xrp1<sup>+/-</sup>*) | *FRT82B, xrp1[m273], RpS3[Plac92]/+* | 25°C |
| 2a | *tub-Gal80[ts]/+; UAS-xrp1/en-Gal4, GstD1-GFP* | 18°C for 8-9 days, 27°C for 24h |
| 2c | *tub-Gal80[ts]/+; UAS-xrp1/en-Gal4, GstD1-GFP* | 18°C for 8-9 days, 27°C for 24h |
| 2e | *tub-Gal80[ts]/+; UAS-xrp1/en-Gal4, GstD1-GFP* | 18°C for 8-9 days, 27°C for 24h |
| 3a | *en-Gal4, GstD1-GFP/+; UAS-GADD34-RNAi/+* | 25°C |
| 3c | *en-Gal4, GstD1-GFP/+; UAS-GADD34-RNAi/+* | 25°C |
| 3e | *en-Gal4, GstD1-GFP/+; UAS-GADD34-RNAi/+* | 25°C |
| 3g | *en-Gal4, GstD1-GFP/+; UAS-GADD34-RNAi/+* | 25°C |
| 3i | *hs-FLP/+; tub>CD2>Gal4, UAS-CD8-GFP/+; tub-Gal80[ts] / +* | 25°C for 3 days, 35 min heat shock, 18°C for 3 days, 29°C for 24h |
| 3j | *hs-FLP/+; tub>CD2>Gal4, UAS-CD8-GFP/+; tub-Gal80[ts] / UAS-GADD34-RNAi* | 25°C for 3 days, 35 min heat shock, 18°C for 3 days, 29°C for 24h |
| 4a | *hs-FLP/+; VDRC[60101]/+; act>CD2>Gal4, UAS-GFP/UAS-GADD34-RNAi* | 25°C for 3 days, 20 min heat shock, 25°C for 3 days |
| 4b | *hs-FLP/+; UAS-xrp1-RNAi/+; act>CD2>Gal4, UAS-GFP/UAS-GADD34-RNAi* | 25°C for 3 days, 20 min heat shock,<br>25°C for 3 days |
| 4d | *en-Gal4, GstD1-GFP/ VDRC[60101]; UAS-GADD34-RNAi/+* | 25°C |
| 4f | *en-Gal4, GstD1-GFP/ UAS-xrp1-RNAi; UAS-GADD34-RNAi/+* | 25°C |
| 4h | *en-Gal4, UAS-GFP/+; FRT82B, xrp1-lacZ/UAS-GADD34-RNAi* | 25°C |
| 4j | *tub-Gal80[ts]/+; en-Gal4, GstD1-GFP/+; FRT82B, xrp1-lacZ/UAS-ATF4* | 18°C for 8-9 days, 29°C for 24h |
| 5a | *hs-FLP/+; VDRC[60101]/+; act>CD2>Gal4, UAS-GFP/UAS-nrf2* | 25°C for 3 days, 20 min heat shock,<br>25°C for 3 days |
| 5b | *hs-FLP/+; UAS-xrp1-RNAi/+; act>CD2>Gal4, UAS-GFP/UAS-nrf2* | 25°C for 3 days, 20 min heat shock,<br>25°C for 3 days |
| 5d | *hs-FLP/+; VDRC[60101]/+; act>CD2>Gal4, UAS-GFP/UAS-nrf2* | 25°C for 3 days, 20 min heat shock,<br>25°C for 3 days |
| 5e | *hs-FLP/+; UAS-irbp18-RNAi/+; act>CD2>Gal4, UAS-GFP/UAS-nrf2* | 25°C for 3 days, 20 min heat shock,<br>25°C for 3 days |
| Supplementary figures |  |  |
| Figure S1a | *en-Gal4, UAS-GFP/ UAS-xrp1-RNAi; FRT82B, xrp1-lacZ/+* | 25°C |
| Figure S1c | *en-Gal4, UAS-GFP/ UAS-irbp18-RNAi; FRT82B, xrp1-lacZ/+* | 25°C |
| Figure S1e | *hs-FLP, UAS-CD8-GFP/+; VDRC[60101]/+; FRT82B, RpS3[Plac92], act>RpS3>Gal4/+* | 25°C for 3 days, 25 min heat shock,<br>25°C for 3 days |
| Figure S1f | *hs-FLP, UAS-CD8-GFP/+; UAS-xrp1-RNAi /+; FRT82B, RpS3[Plac92], act>RpS3>Gal4/+* | 25°C for 3 days, 25 min heat shock,<br>25°C for 3 days |
| Figure S1h | *hs-FLP, UAS-CD8-GFP/+; VDRC[60100]/+; FRT82B, RpS3[Plac92], act>RpS3>Gal4/+* | 25°C for 3 days, 25 min heat shock,<br>25°C for 3 days |
| Figure S1i | *hs-FLP, UAS-CD8-GFP/+; UAS-irbp18-RNAi /+; FRT82B, RpS3[Plac92], act>RpS3>Gal4/+* | 25°C for 3 days, 25 min heat shock,<br>25°C for 3 days |
| Figure S1k | *hs-FLP; VDRC[60101]/+; act>CD2>Gal4, UAS-GFP/UAS-mahj-RNAi* | 25°C for 3 days, 20 min heat shock,<br>25°C for 3 days |
| Figure S1l | *hs-FLP; UAS-xrp1-RNAi/+; act>CD2>Gal4, UAS-GFP/UAS-mahj-RNAi* | 25°C for 3 days, 20 min heat shock,<br>25°C for 3 days |
| Figure S2a | *en-Gal4, GstD1-GFP/+; UAS-GADD34-RNAi/+* | 25°C |
| Figure S2d | *hs-FLP/+; VDRC[60101]/+; act>CD2>Gal4, UAS-GFP/UAS-GADD34-RNAi* | 25°C for 3 days, 20 min heat shock,<br>25°C for 3 days |
| Figure S2e | *hs-FLP/+; VDRC[60101]/+; act>CD2>Gal4, UAS-GFP/UAS-GADD34-RNAi* | 25°C for 3 days, 20 min heat shock,<br>25°C for 3 days |
| Figure S2g | *hs-FLP/+; tub>CD2>Gal4, UAS-CD8-GFP/+; tub-Gal80[ts] / UAS-GADD34-RNAi* | 25°C for 3 days, 35 min heat shock,<br>18°C for 3 days, 29°C for 24h |

### Proteomics

Sample preparation and Tandem Mass Tag (TMT) mass spectrometry were performed as described in (Baumgartner et al., 2021).

### Image acquisition and processing

Images were acquired using a Leica SP8 confocal microscope with a 40x 1.3 NA P Apo Oil objective. Wing discs were imaged as z-stacks with each section corresponding to 1μm. Images were processed using Photoshop (Adobe Photoshop 2020) and Fiji (Version 2).

### Quantifications

Clonal areas, cell death quantifications and fluorescence intensity quantifications were carried out using custom built Fiji scripts. All analysis focused on the pouch region of the wing disc. For clone area measurements, the percentage of the volume of the pouch occupied by clones was determined. For cell death quantifications the clone border is defined as any cell within a 2 cell-range of the clone boundary. Cell death measurements were normalized to the respective area of the clone border or clone center as measured in Fiji. For all scatter plots the horizontal line represents the median.

### Statistics and reproducibility

All data represented by the scatter plots including details of the specific statistical test used for each experiment are shown in the Statistics Source Data table. Statistics were performed using GraphPad Prism (Prism 8). Univariate statistics were used to determine P-values. Parametric tests were used where assumptions of normality were met, otherwise non-parametric tests were used. The parametric test used was the Student’s T-Test and the non-parametric tests used were the Mann Whitney U-test for non-paired data, and the Wilcoxon matched-pairs signed rank test for paired data. P-value corrections for multiple comparisons were not considered due to the low number of comparisons. For experiments comparing across wing discs a minimum of three biological repeats were performed. For experiments with an internal control, a minimum of two biological repeats were performed. Experiments performed to validate reagents (e.g., testing efficacy of RNAi lines) were carried out at least once.

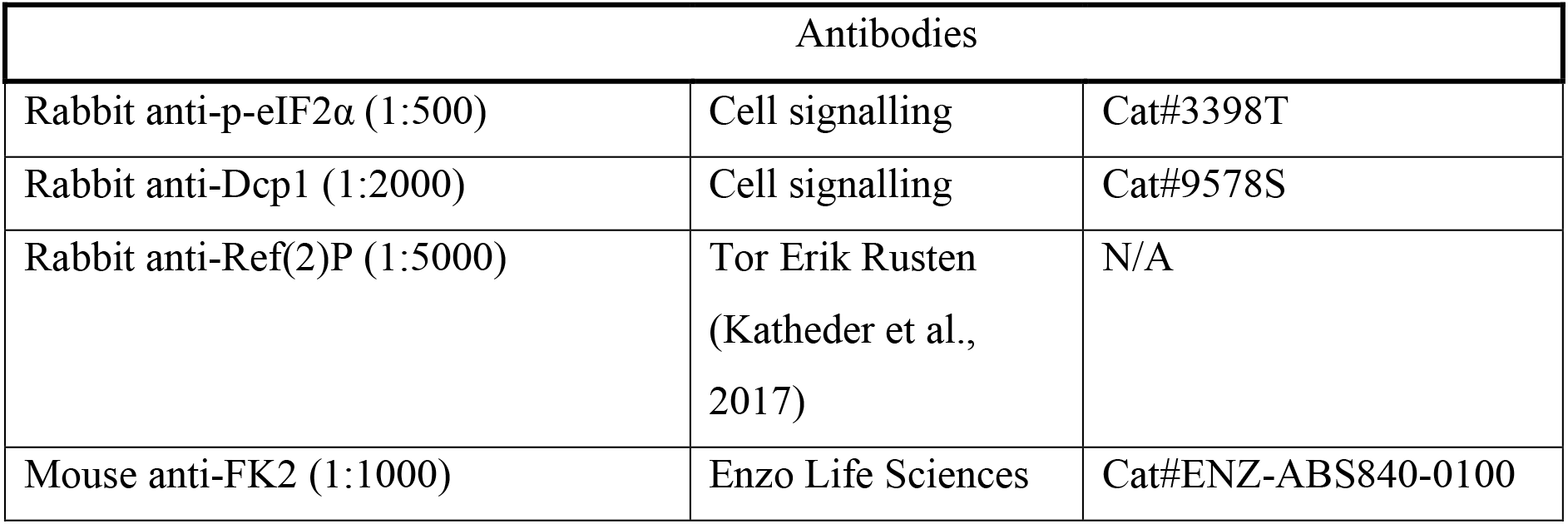
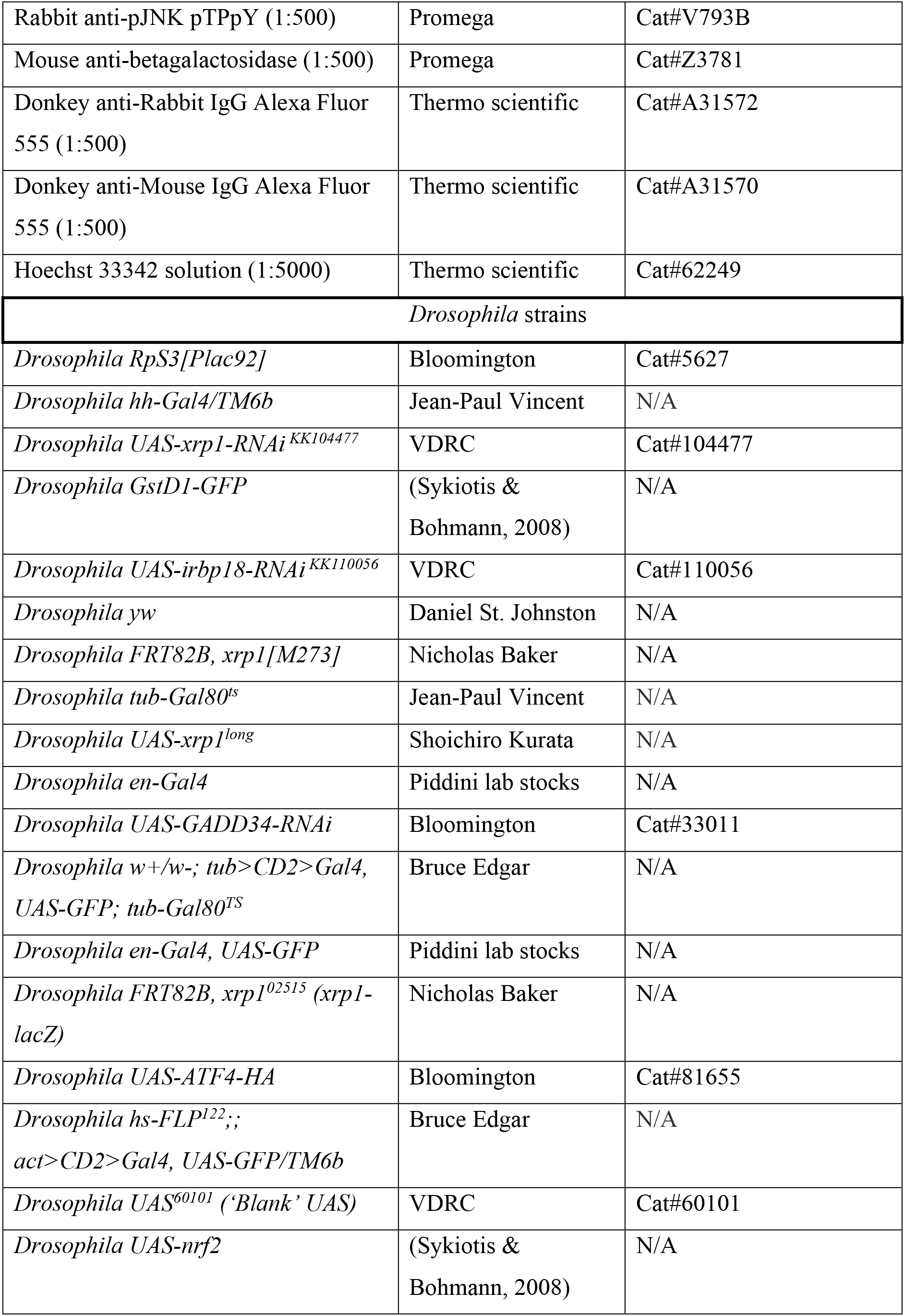
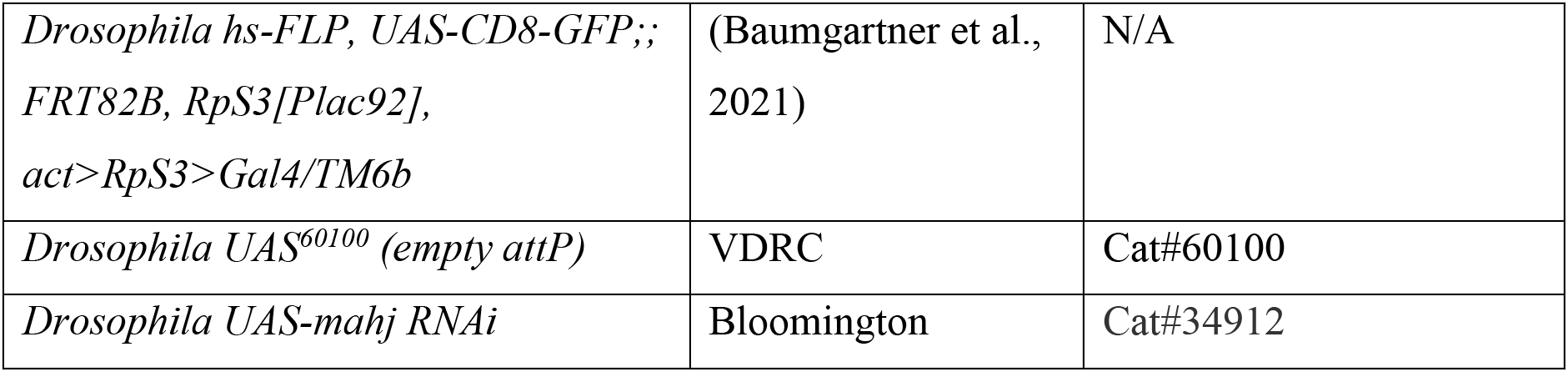
Key Resources Table.

## Notes

### Competing Interest Statement

The authors have declared no competing interest.

